# A cell size threshold triggers commitment to stomatal fate in *Arabidopsis*

**DOI:** 10.1101/2022.10.09.510391

**Authors:** Yan Gong, Renee Dale, Hannah F. Fung, Gabriel O. Amador, Margot E. Smit, Dominique C. Bergmann

## Abstract

How flexible developmental programs integrate information from internal and external factors to modulate stem cell behavior is a fundamental question in developmental biology. Cells of the *Arabidopsis* stomatal lineage modify the balance of stem cell proliferation and differentiation to adjust the size and cell type composition of mature leaves. Here, we report that meristemoids, one type of stomatal lineage stem cell, trigger the transition from asymmetric self-renewing divisions to commitment and terminal differentiation by crossing a critical cell size threshold. Through computational simulation, we demonstrate that this cell size-mediated transition allows robust, yet flexible termination of stem cell proliferation and we observe adjustments in the number of divisions before the differentiation threshold under several genetic manipulations. We experimentally evaluate several mechanisms for cell size sensing, and our data suggest that cell size is sensed via a chromatin ruler acting in the nucleus.

## Introduction

During development, stem cells balance competing needs for proliferation and differentiation to control the final size and cellular composition of tissues. In systems that experience unpredictable or variable conditions, such as the gut epithelium (O’Brien et al., 2011) and muscle satellite cell (Liu et al., 2012; Motohashi & Asakura, 2014) lineages, a flexible developmental program requires that this balance be dynamically altered in response to internal or external cues. The *Arabidopsis* stomatal lineage is a model for the study of such flexible developmental programs (Lee & Bergmann, 2019). During leaf growth, stem cells of the stomatal lineage—meristemoids and stomatal lineage ground cells (SLGCs)—undergo a variable number of self-renewing asymmetric cell divisions (ACDs) before committing to terminal differentiation as stomata or pavement cells, respectively (Figure 1A). Notably, both meristemoid and SLGC proliferation can be tuned by hormonal, nutrient and environmental inputs (Balcerowicz et al., 2014; Engineer et al., 2014; Gong, Alassimone, Varnau, et al., 2021; Han et al., 2020; Lau et al., 2018; Le et al., 2014; Lee et al., 2017; Vaten et al., 2018). This flexible developmental program is deeply conserved across land plants and functions to optimize mature leaf physiology to an individual’s unique local environment (Chater et al., 2017; McKown & Bergmann, 2020).

**Figure 1.**
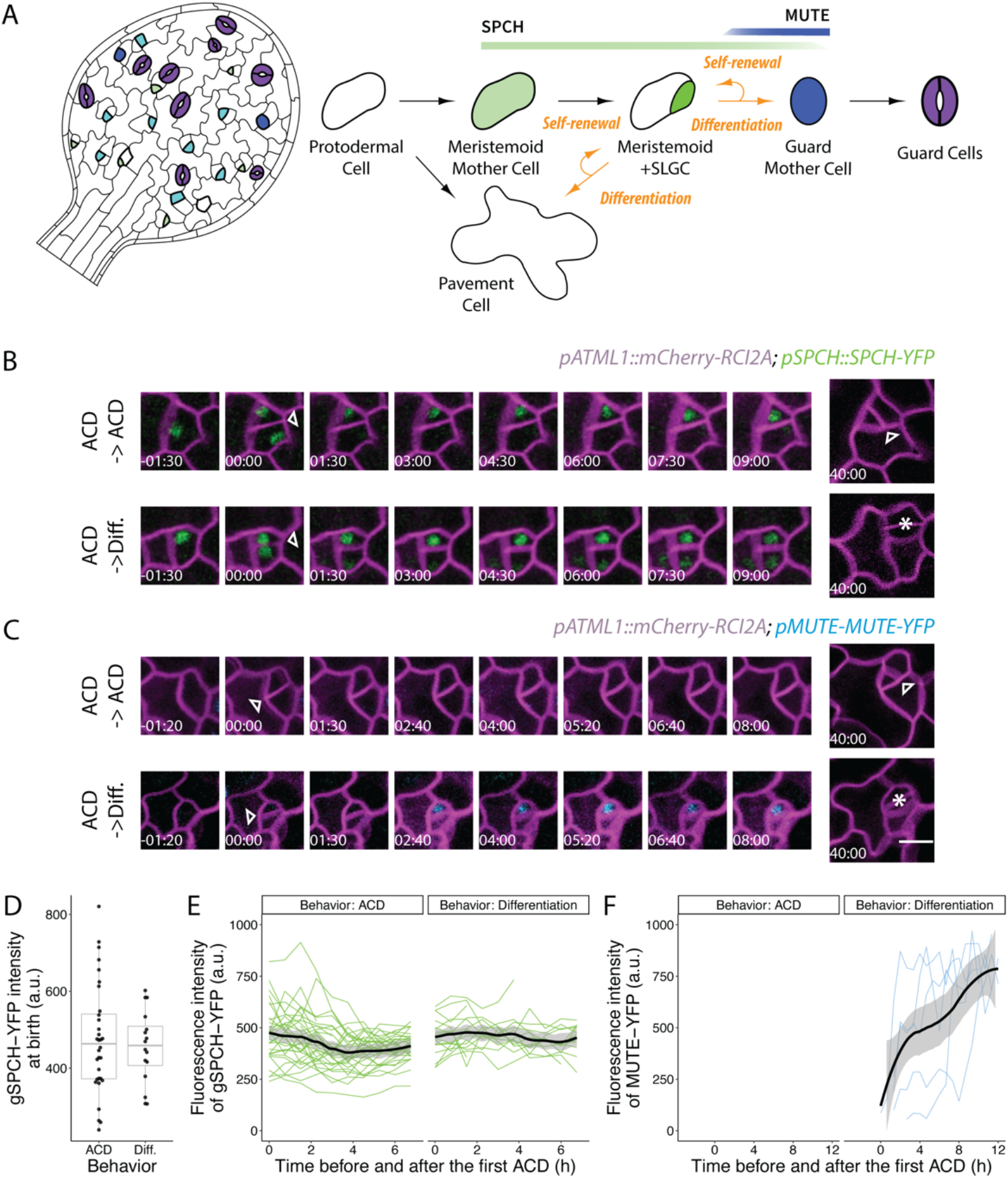
Transitions from self-renewal to differentiation in stomatal lineage meristemoids are not predicted by expression of the transcription factors SPCH or MUTE. (A) Cartoon of stomatal lineage cell and lineage trajectory (right) and their arrangement on the leaf epidermis (left). Protodermal precursor cells enter the lineage and undergo asymmetric “entry” divisions producing a smaller meristemoid (green) and a larger stomatal lineage ground cell (SLGC, white) daughter. Both meristemoids and SLGCs can undergo additional self-renewing asymmetric cell divisions (ACDs). Alternatively, the SLGC can undergo terminal differentiation into a pavement cell and the meristemoid can transition into a guard mother cell (GMC, blue), undergo one round of symmetric division, and produce paired guard cells (purple). At any given time during leaf development, dispersed stomatal lineages are actively initiating, dividing, and differentiating. (B-C) Time-lapse analysis of the dynamics of *pSPCH∷SPCH-YFP* (B, green) and *pMUTE∷MUTE-YFP* (C, blue) reporters during ACDs in 3 dpg cotyledons followed by lineage tracing. SPCH and MUTE reporter were imaged every 40 mins for 16 hours, returned to ½ MS plates, and re-imaged to capture the division and fates of the meristemoids daughter from ACDs. Two examples are shown for each reporter where the daughter meristemoid either undergoes another ACD to renew itself or a GMC division to differentiate into a stoma. 00:00 (hours: minutes) marks cell plate formation. Cell outlines are visualized by plasma membrane reporter *pATML1∷RCI2A-mCherry* (magenta). Arrowheads and asterisks indicate asymmetric meristemoid and symmetric GMC divisions, respectively. (D) Quantification of SPCH-YFP reporter levels at birth during ACDs when daughter meristemoids either undergo additional ACDs or differentiate and become stomata. (E-F) Quantification of SPCH-YFP reporter (E) or MUTE-YFP reporter (F) levels at birth and dynamics during ACDs with either behavior. Intensity of SPCH-YFP and MUTE-YFP reporters is tracked with TrackMate (Tinevez et al., 2017) during 5 - 34 ACDs (thin colored lines) for each group, and the respective trend per each condition with 0.95 confidence interval is indicated as the thick line with gray band. Scale bars, 10 μm.

A group of closely related basic helix-loop-helix (bHLH) transcriptional factors control stem cell behavior in the stomatal lineage (Kanaoka et al., 2008; MacAlister et al., 2007; Ohashi-Ito & Bergmann, 2006; Pillitteri et al., 2007). Among these transcriptional factors, SPEECHLESS (SPCH) initiates the lineage and is required for continued proliferation of both meristemoids and SLGCs (Lopez-Anido et al., 2021; MacAlister et al., 2007), while MUTE, a direct target of SPCH, is activated in meristemoids after one to three asymmetric divisions and triggers terminal differentiation towards stomatal fate (Han et al., 2018; Pillitteri et al., 2007). Whereas SPCH is necessary for eventual activation of MUTE, the factors that determine the timing of the SPCH to MUTE transition, and thus of terminal differentiation into stomata, are currently unknown.

Here, we show that commitment to stomatal fate is triggered by crossing a cell size threshold. Repeated asymmetric division of meristemoids decreases cell size until the critical threshold is reached. Genetic manipulations of initial meristemoid size and division asymmetry directly affect the rate of decay of stem cell size and thus the timing of the switch between self-renewing proliferation and terminal commitment. Modeling stem cell behavior shows that imposing a cell size threshold for differentiation is sufficient to accurately predict the number of asymmetric divisions stem cells undergo before transitioning and suggests a mechanism by which cell size may integrate environmental information to control the rate of amplifying division in developing leaves. Molecular mechanisms known to be available to make decisions based on cell size rely on scaling relationships, either in protein levels (Xie et al., 2022) or related to geometry (Hubatsch et al., 2019); our experimental evaluation of candidate molecules and mechanisms in the stomatal lineage suggests that size is actually sensed in the nucleus and uses chromatin as a ruler.

## Results

### SPCH and MUTE do not drive the transition from proliferation to differentiation in meristemoids

Multiple external and internal factors have been reported to affect the balance of stem cell proliferation and differentiation in the *Arabidopsis* stomatal lineage. Some of these factors influence the lineage by modulating SPCH levels (Lau et al., 2018; Lee et al., 2017; Vaten et al., 2018), prompting us to investigate whether SPCH levels alone are sufficient to predict whether a meristemoid will proliferate or differentiate.

To monitor SPCH protein dynamics, we captured time-lapse images of the epidermis of a 3 day post germination (dpg) cotyledon expressing a *SPCH* translational reporter (*spch-3 SPCHp∷gSPCH-YFP*). We found that SPCH levels after birth were poor predictors of future meristemoid behavior (Figure 1B, D, E). Indeed, meristemoids expressed high levels of SPCH after birth, irrespective of fate (Figure 1D-E, Figure S1). In contrast, MUTE protein levels were highly correlated with meristemoid behavior, appearing exclusively in meristemoids that would differentiate several hours later (Figure 1C, F). This is consistent with MUTE’s role in establishing GMC identity (Han et al., 2018), but also suggests that *MUTE* expression is a consequence, rather than a cause, of the decision to differentiate.

With the resolution made possible by single cell RNA sequencing (scRNA-seq), we also investigated whether sub-classes of meristemoids could be distinguished by considering SPCH in combination with other suites of genes. Although several independent scRNA-seq studies exquisitely resolved a unidirectional trajectory from *MUTE* expression to stomatal guard cell differentiation (Lopez-Anido et al., 2021; Zhang et al., 2021), none identified distinct groups of SPCH-expressing cells that could be mapped on to self-renewing or differentiating behaviors.

### A cell size threshold for the M-GMC transition

To identify factors that might influence the decision to differentiate, we surveyed the literature for correlates of meristemoid behavior. Notably, Robinson et al. (2011) reported that meristemoid size declines with successive asymmetric divisions, due to a combination of division asymmetry and minimal growth between divisions, and we confirmed these findings in our data (Figure S1A-B). This observation, coupled with the fact that most meristemoids divide at least once before differentiating (Geisler et al., 2000), implies that cells that divide are larger than cells that differentiate.

To investigate whether size predicts meristemoid behavior, we quantified the birth sizes of meristemoids and recorded their subsequent behaviors (Figure 2A-B). Birth size was operationalized as the cross-sectional area at birth, which correlates strongly with volume (Robinson et al., 2018; Willis et al., 2016). Upon fitting the data to a logistic regression, we found that birth size was highly predictive of meristemoid behavior (average classification accuracy: 91% ± 4.4%, see Methods): smaller meristemoids were likely to differentiate, whereas larger meristemoids were likely to self-renew (Figure 2C). We defined the transition size as the size at which half of the meristemoids were expected to self-renew (32 μm^2^ in 3 dpg wild-type cotyledons).

**Figure 2.**
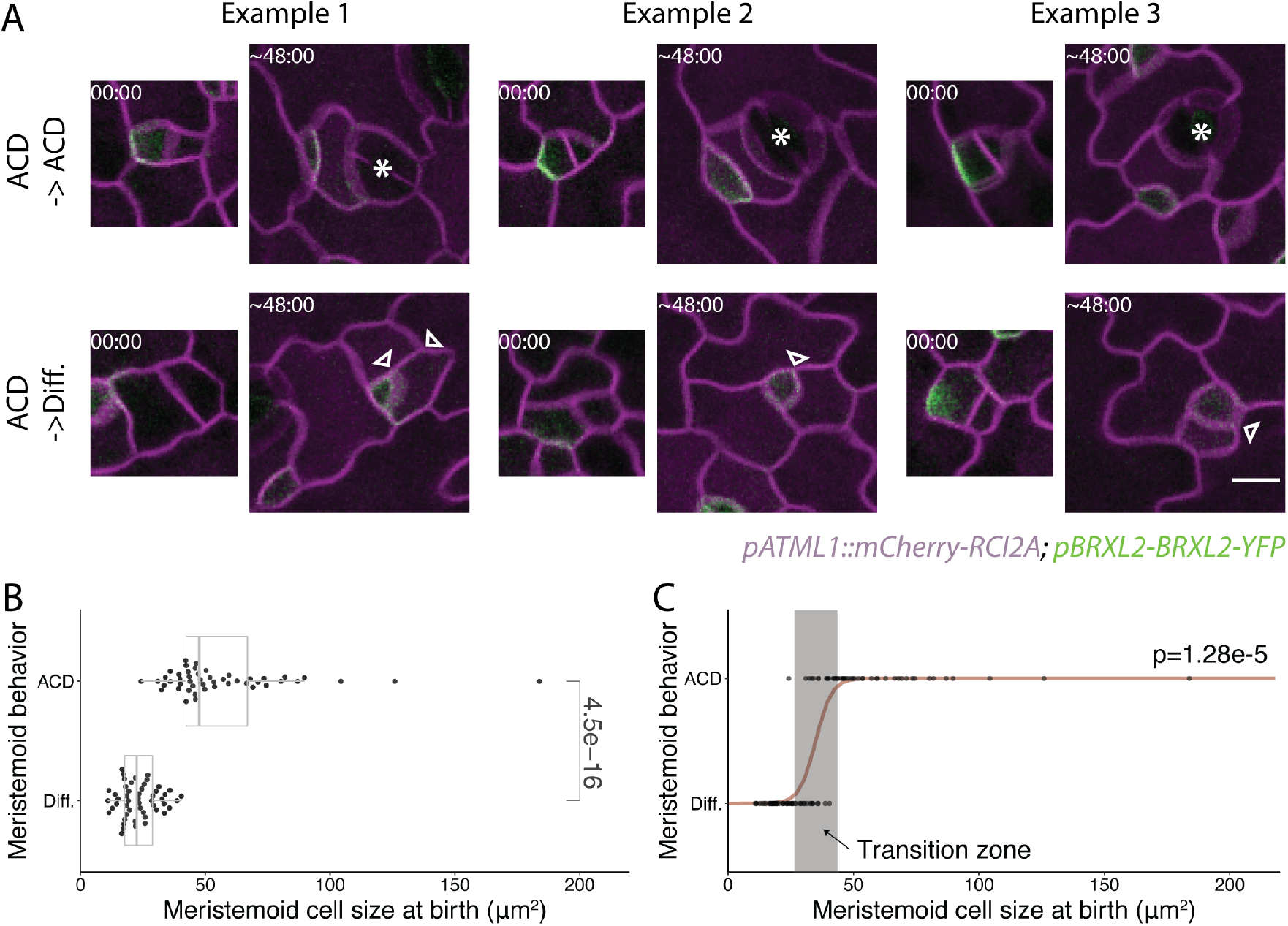
The M-GMC transition of the stomatal lineage is correlated with small meristemoid cell size at birth. Confocal images of meristemoids at birth and the subsequent division behaviors of these meristemoids captured by lineage tracing in 3 dpg cotyledons. Meristemoids are divided into two groups based on their subsequent division behaviors: those that undergo additional ACDs or those that differentiate and become stomata. Three examples are shown for each group. Cell outlines are visualized by plasma membrane reporter *pATML1∷RCI2A-mCherry* (magenta). The cell polarity reporter *pBRXL2∷BRXL2-YFP* (green) was include to define cell division type. 00:00 (hours: minutes) marks cell plate formation. Arrowheads and asterisks indicate ACDs of meristemoids and GMC divisions, respectively. (B) Comparison of areal cell size at birth between meristemoids that acquire different fates (n= 50 cells/group). (C) Logistic regression of meristemoid behaviors based on their cell size at birth. The cell size of each meristemoid is shown as a single dot and the computed regression model is shown in dark red. The predicted transition zone where the meristemoid is predicted to have 10% to 90% probability of undergoing another ACD is shown in a gray box. The p-value in (B) is calculated by Mann-Whitney test, and the p-value in (C) is calculated by the glm.fit function with a binomial model in R (R Core Team, 2020). Scale bars, 10 μm.

To determine whether this relationship between size and behavior is a general feature of meristemoid differentiation, we examined the distantly related eudicot tomato (*Solanum lycopersicum*). We observed a similar size bias between self-renewing and differentiating meristemoids (Figure S3), suggesting size-restricted stomatal differentiation may be a conserved feature of leaf development across land plants. In tomato, meristemoids can also undergo non-stomatal differentiation into pavement cells (Figure S3A, (Nir et al., 2022)). As there is no size bias in this differentiation pathway (Figure S3B), size control appears to be a specific feature of stomatal differentiation.

### A lineage decision tree model with size as the sole determinant is sufficient to explain meristemoid behaviors

Our data suggest there is a size threshold below which meristemoids are likely to differentiate. To test whether a size threshold alone is sufficient to explain observed meristemoid behaviors, we specified a stochastic and asynchronous rule-based lineage decision tree model, with size as the sole determinant of meristemoid differentiation (Figure 3A, left). In this model, 10,000 meristemoids enter the stomatal lineage, each with a size randomly drawn from an empirically derived starting size distribution. These meristemoids divide asymmetrically, producing a meristemoid and an SLGC. The SLGCs are removed from the model, while the meristemoid undergoes a size-guided differentiation program in which smaller meristemoids have a higher chance of differentiating and exiting the model. The remaining undifferentiated meristemoids continue to grow, asymmetrically divide, and/or differentiate until all meristemoids have differentiated. The following parameters were estimated from cotyledons imaged from 3 to 5 dpg: meristemoid birth sizes, division asymmetry, the probability of meristemoid differentiation given its birth size, and cell cycle duration (Figure 3A, right).

**Figure 3.**
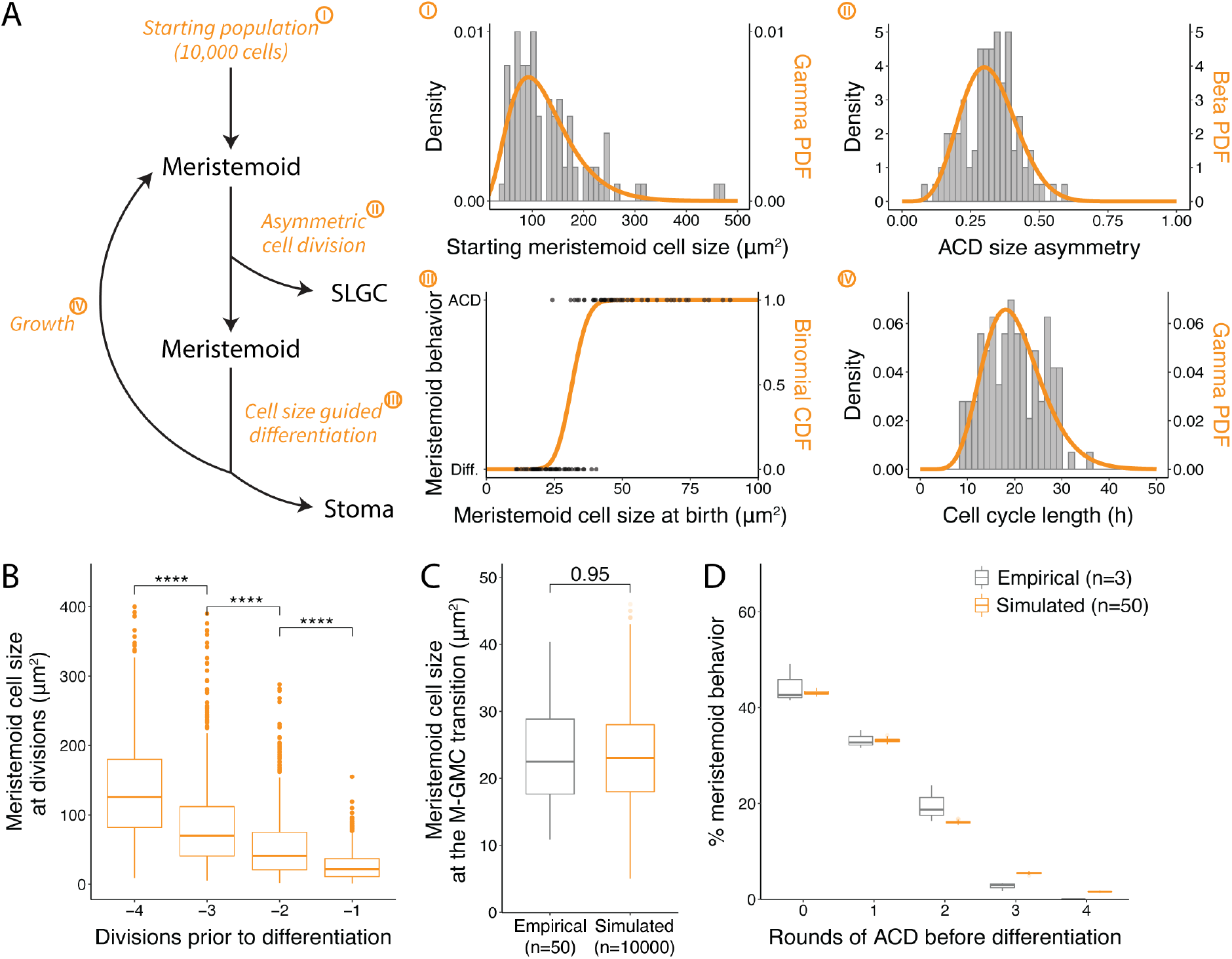
Cell size guided M-GMC transition is sufficient to explain the self-renewal and differentiation behavior of meristemoids *in silico*. (A) The workflow of the meristemoid division tree model (left) and its key parameters (I-IV, right). 10,000 meristemoids are randomly drawn from a starting population, gamma distribution [4, 33.3] (I). These meristemoid then divide asymmetrically, each with a size asymmetry (size of the meristemoid daughter divided by the size of the mother cells) drawn from a beta distribution [6.8, 14.6] (II). The newly formed meristemoid daughter was then passed on to a cell-size-guided differentiation model while the SLGC was discarded. In the cell size guided differentiation model, each meristemoid differentiates with some probability based on current size using the binomial cumulative distribution function (CDF) [current size, 100,0.32] (III). A cell of 32 square microns will divide 50% of the time. Differentiated meristemoids leave the model while the rest grow with 3% growth rate (per hour) with a cell cycle length drawn from a gamma distribution [10, 20] (IV). After growth, these meristemoids are then looped back to divide asymmetrically again and pass through the rest steps of the model until all 10000 meristemoids differentiate and leave the model. (I) Histogram of the measured starting meristemoid cell size (gray, n=132 cells) and the fitted gamma distribution probability density function (PDF) (orange). (II) Histogram of the measured ACD size asymmetry (gray, n=98 cells) and the fitted beta distribution PDF (orange). (III) Dot plot of the measured meristemoid cell size at birth separated by their fates (gray, n=98 cells) and the fitted binomial distribution CDF (orange). (IV) Histogram of the measured cell cycle length for amplifying division (gray, n=112 cells) and the fitted gamma distribution PDF (orange). (B-D) Outputs of the meristemoid division tree model. (B) Computed meristemoid cell size at ACDs before differentiation (n=913 cells). (C) Comparison of the empirical (n=50 cells) and simulated (n=10000 cells) meristemoid cell size at birth before undergoing M-GMC transition. (D) Comparison of the empirical (n>300 cells/replicate/genotype) and simulated (n=10000 cells/replicate/genotype) meristemoid division-differentiation behavior. Empirical data are taken from lineage tracing experiments where each individual behavior of the abaxial cotyledon of corresponding genotypes are tracked for their cell divisions and differentiation behavior from 3 dpg to 5dpg (Gong, Alassimone, et al., 2021). All p-values are calculated by Mann-Whitney test.

The model outputs included meristemoid size before and after each round of ACD, the number of meristemoids that differentiated after each round of ACD, and meristemoid size at differentiation. We compared these outputs to empirical data from our time-lapse analyses (Figure 3B-D) and to previously reported studies (Robinson et al., 2011). Consistent with Robinson et al. (2011), meristemoid size declined with each successive division: meristemoid daughters were, on average, 33% smaller than their parents (Figure 3B), consistent with empirical estimates (Figure S2B), Additionally, simulated meristemoids differentiated at an average size of 22 μm^2^, which is similar to the empirical estimate of 23.2 μm^2^ (Figure 3C). We tested the sensitivity of the model to input parameters and observed that initial size, growth rate and size of differentiation have strong effects on the number of divisions before differentiation (Figure S4). These simulations show that a differentiation program accounting for birth size alone is sufficient to recapitulate the division and differentiation behaviors of wild-type meristemoids (Figure 3D, (Gong, Alassimone, Varnau, et al., 2021).

### Manipulating cell size affects the proliferative capacity of meristemoids

Previously, we identified *CONSTITUTIVE TRIPLE RESPONSE1*(*CTR1)* as a positive regulator of meristemoid proliferation (Gong, Alassimone, Varnau, et al., 2021), although the detailed mechanism underlying this effect was unknown. Notably, Vaseva et al. (2018) reported that epidermal cell expansion was inhibited in the *ctr1* mutant, leading us to investigate whether the lack of cell expansion and consequently, a reduction in cell size, could trigger premature differentiation. While *ctr1* epidermal cells were the same size as wild-type cells at germination (0 dpg; Figure 4A, B), *ctr1* meristemoids were significantly smaller at 4 dpg (Figure 4C), due to lower expansion rates (~2% vs. ~3% per hour). *ctr1* meristemoids underwent fewer ACDs before differentiating (Figure 3D-E), suggesting that meristemoids actively sense their size and adjust their behavior accordingly.

**Figure 4.**
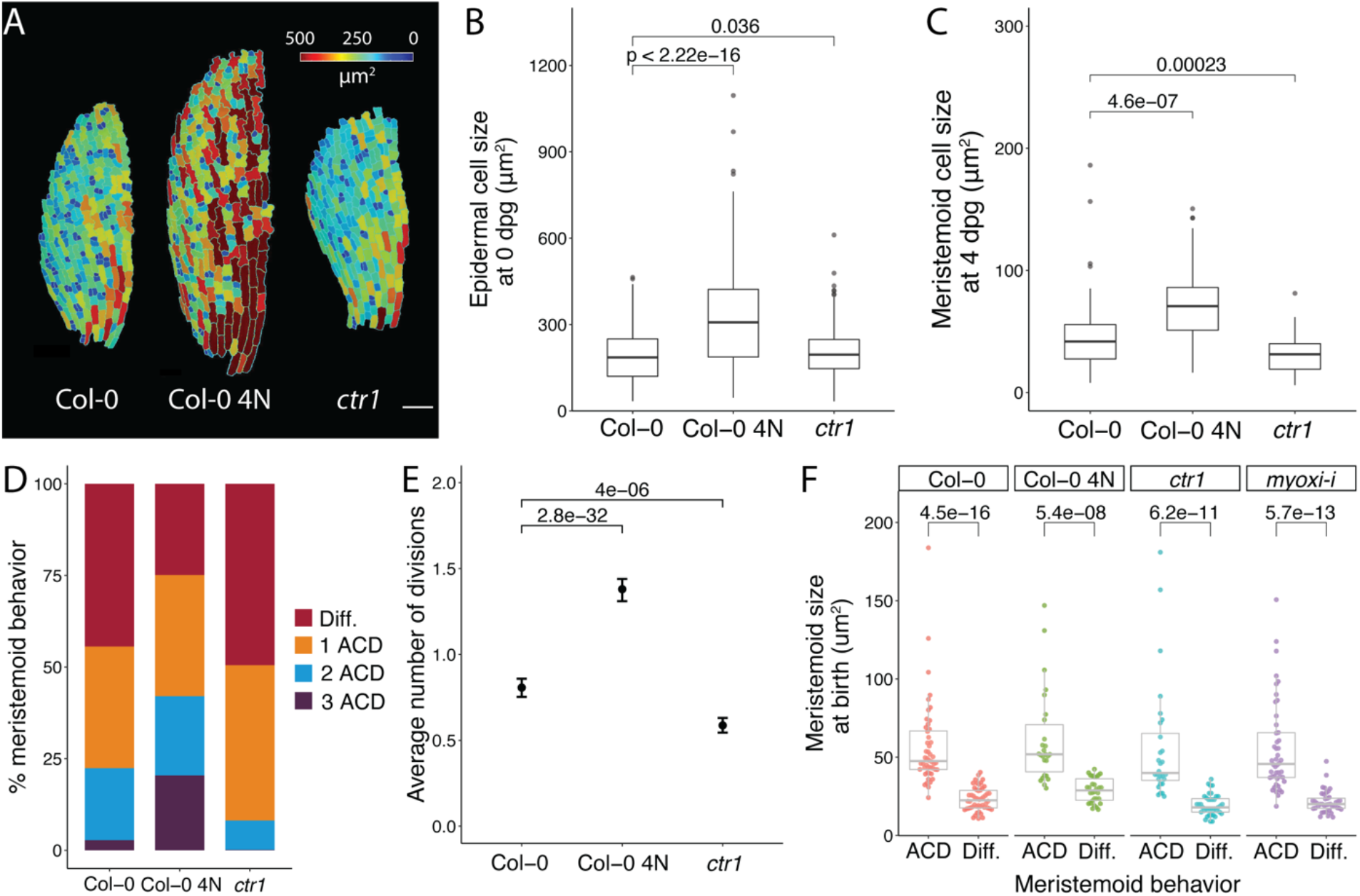
Alteration of the meristemoid size and ACD asymmetry affects number of successive meristemoid ACDs but not the cell size of the M-GMC transition. (A-C) Comparison of cell size for leaf epidermal cells in wild type Col-0, tetraploid Col-0 (Col-0 4N), and the *ctr1* mutant at different stages of development. (A) False-colored confocal images of the abaxial epidermis of 0 dpg cotyledons from different genetic backgrounds. mPS-PI staining images of half of the cotyledons were segmented and false-colored based on cell size in MorphoGraphX (Barbier de Reuille et al., 2015). (B) Cell size distribution of epidermal cells in Col-0, Col-0 4N, and *ctr1* cotyledons at 0 dpg (n>500 cells/genotype). (C) Cell size distribution of meristemoids in Col-0, Col-0 4N, and *ctr1* cotyledons at 4 dpg (n>50 cells/genotype). Meristemoids were selected from confocal images of 4 dpg cotyledons (labelled with the plasma membrane reporter *pATML1∷RCI2A-mCherry*) with their cell size (surface area) measured in FIJI (Schindelin et al., 2012). (D-E) Division-differentiation behavior of meristemoid population from 3 dpg to 5dpg (n>300 cells/genotype), shown as the distribution (D) or as its mean, counting differentiation as zero (E). Data of Col-0 are adapted from Gong, Alassimone, et al. (2021). (F) Comparison of cell size at birth between meristemoids that acquire different fates in Col-0, Col-0 4N, *ctr1* and the *myoxi-i* mutant (n> 50 cells/genotype). The data of Col-0 are taken from Figure 2B. All p-values are calculated by Mann-Whitney test. Scale bars, 10 μm.

If decreasing cell size causes premature differentiation, then increasing cell size should increase proliferative capacity. Because genome size often correlates with cell size in *Arabidopsis* (Li et al., 2012) we examined whether an induced tetraploid line (Robinson et al., 2018) had larger stomatal lineage cells. Quantification of cell size from still confocal images revealed that both leaf epidermal cells (Figure 4A, B) and meristemoids (Figure 4C) were significantly larger in the tetraploid. This increase in meristemoid size was accompanied by an increase in the number of divisions prior to differentiation (Figure 4D-E), lending further support to our hypothesis that cell size, or some property derived directly from size, drives stomatal differentiation. Importantly, the relationship between size and differentiation is robust to genetic perturbation of cell size, as in *ctr1* or tetraploids, or when division asymmetry is reduced (as in *myosin-xi*, (Muroyama et al., 2020) such that size remained predictive of meristemoid behavior (Figure 4F).

As stomatal size is tightly linked to transpirative capacity (Franks & Beerling, 2009), we hypothesized that size-gated differentiation of stomatal precursors may set the mature size of stomata. However, in a transgenic line where meristemoids differentiate at unusually small sizes, mature stomata are not, on average, smaller nor do we observe especially small stomata (Figure S5). These data are consistent with additional layers of size control regulating guard cell growth during their twenty-fold expansion from a birth size of ~25 μm^2^ to a stomatal size of ~500 μm**^2^** at maturity.

### Cells measure the ratio of chromatin to nuclear size

Several stem cell lineages sense cell size to inform choices about cell cycle progression, division and differentiation. For example, in the *C. elegans* embryo, the P lineage undergoes four consecutive asymmetric divisions before switching to symmetric division at a threshold size where cells are too small to sustain PAR polarity (Hubatsch et al., 2019). We tested whether a similar mechanism may sense size in the leaf epidermis. As in *C. elegans*, polar crescents occupy a larger fraction of the cortex in small cells than in large cells, approaching ~40% of the total circumference in the small meristemoids fated to differentiate (Figure S6A). Expanding the size of the crescent, by expressing a pBASL∷BRX-CFP transgene, would be expected to cause cells to differentiate at a larger size; however, we find that cells actually differentiate at slightly smaller sizes upon this manipulation (Figure S6B-E). Thus, if cell polarity regulates size-dependent differentiation, the mechanism is not the same as that described for the *C. elegans* P lineage.

A polarity-independent mechanism for cell size sensing was recently described in the shoot apical meristem, where cells inherit a fixed amount of the cell cycle progression inhibitor KRP4 in proportion to their ploidy and read out the concentration of KRP4 in nuclei of varying sizes (D’Ario et al., 2021). As nuclear size is strongly correlated to overall cell size, this “chromatin ruler” model allows cells to delay cell cycle progression until growing to a target size. To determine whether meristemoids are sensing their cell or nuclear size, we sought a genetic manipulation that would change nuclear size without affecting cell size. The loss-of-function *crwn1-1* allele (Dittmer et al., 2007) satisfies these criteria. *CRWN1* encodes a plant lamin-like protein that is involved in building the meshwork structure of the nuclear lamina (Sakamoto et al., 2020). Plants homozygous for the *crwn1-1* allele harbor smaller nuclei at birth (*t*(154) = 6.93, *p* = 1.08e-10; Figure 5A), but their meristemoids remain wild-type-sized (W = 3069.5, *Z* = −0.31, *p* = 0.76; Figure 5B). Nuclear and cell areas remain positively correlated in *crwn1* mutants (*r* = 0.63, *t*(48) = 5.6, *p* = 9.29e-7; Figure S7). The loss of *CRWN1* is not associated with major changes in gene expression or chromatin accessibility (Hu et al., 2019).

**Figure 5.**
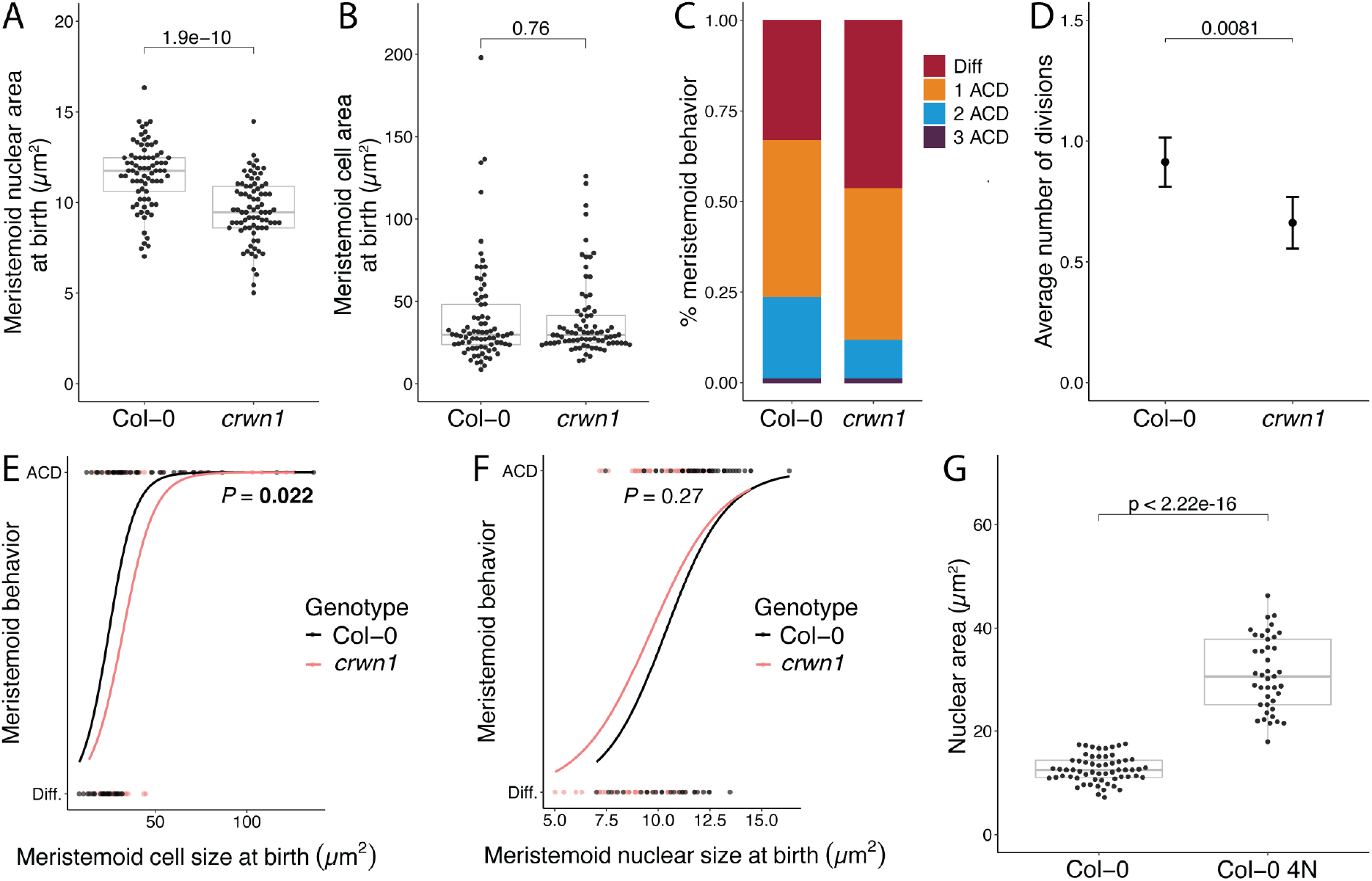
Cell size is sensed via chromatin content in the nucleoplasm. (A-B) Comparisons of nuclear area (A) and cell area for wild-type Col-0 and *crwn1* plants. (C-D) Comparison of the rate of amplifying division for Col-0 and *crwn1* plant, as the distribution (C) or its mean (D). (E-F) Logistic regressions of cell area (E) or nuclear area (F) against meristemoid behavior, showing that relative to Col-0 meristemoids, *crwn1* meristemoids transition to differentiation at the same nuclear size, but a different overall cell size. (G) Comparison of nuclear areas of differentiating meristemoids in diploid and tetraploid Col-0, showing that chromatin content influences the transition size. All p-values are calculated by Mann-Whitney test, except in E-F, where a t.test was performed on the outputs from dose.p (see methods).

If cells were sensing overall size, then *crwn1* meristemoids should proliferate at similar rates as wild-type meristemoids (quantified as the number of amplifying divisions per cell). If, however, cells were sensing nuclear size, then *crwn1* meristemoids should proliferate at lower rates. In other words, they should differentiate at larger cell sizes, but similar nuclear sizes. Lineage tracing experiments of *crwn1* cotyledons (comprising 3 and 5 dpg time points) revealed a significant decrease in the number of amplifying divisions per meristemoid (mixed effects model: *β* = 0.25 ± 0.060, *p* = 0.0019; Figure 5C, D). Through time-lapse imaging, we found that *crwn1* meristemoids differentiate at larger cell sizes (*t*(150) = 2.32, *p* = 0.022; Figure 5E), but similar nuclear sizes (*t*(147) = 1.10, *p* = 0.27; Figure 5F). These data are consistent with a nuclear size sensor.

If the size sensor is nuclear, does it scale with DNA content, as in the shoot apical meristem? To address this question, we re-visited the induced tetraploid line. If size sensing were independent of genome size, then tetraploid meristemoids should transition at similar nuclear sizes as diploid meristemoids, but if the size sensor were scaled to genome size, then tetraploid meristemoids should transition at larger nuclear sizes. Upon quantifying the nuclear areas of differentiating meristemoids, we found that tetraploid meristemoids transition at much larger nuclear sizes than diploid meristemoids (*t*(47.2) = −15.9, *p* < 2.2e-16; Figure 5G), suggesting the nuclear size sensor scales with genome size. Taken together, these data show that meristemoids sense size through a chromatin ruler, raising the question of whether KRP4 could be the size sensor in our system. While the concentration of KRP4 could likewise be elevated in small meristemoids, KRP4 is unlikely to trigger stomatal differentiation directly, given its function as a cell cycle inhibitor and the observation that both meristemoids and GMCs continue to divide.

## Discussion

In this study, we show that commitment to stomatal fate is triggered when meristemoid size falls below a critical threshold. Genetic manipulations of meristemoid size are accompanied by changes in proliferative capacity, indicating that meristemoids actively sense their size, likely through a chromatin ruler.

We draw inspiration from previous work on homeostatic populations of stem cells that use size to control the timing of division (D’Ario et al., 2021; Xie & Skotheim, 2020), often via dilution of a cell cycle progression inhibitor. In contrast to these models, however, meristemoids use size to control the timing of differentiation, not division, and do not display long-term size homeostasis, instead shrinking markedly over successive divisions to reach the differentiation threshold. This system is consistent with a model in which a differentiation factor becomes concentrated over successive divisions. While the identity of this factor is unknown, our work has uncovered several of its features. Through the *crwn1* and tetraploid analyses, we propose that size is sensed through a nuclear factor that scales with, but is not necessarily bound to, chromatin. As meristemoid size declines, the factor becomes sufficiently concentrated and activates the expression of guard mother cell-specific genes, including *MUTE*, to drive terminal differentiation.

Meristemoid shrinkage requires uncoupling cell growth from division. In other shrinking stem cell lineages, such as *Drosophila* type II neuroblasts, shrinkage requires extensive metabolic remodeling during asymmetric division (Homem et al., 2014), including changes in the expression of core oxidative phosphorylation enzymes. It is tempting to speculate that the slow growth rates observed in meristemoids are caused by similar metabolic remodeling. Notably, a recent report (Shi et al., 2022) showed that meristemoids have elevated levels of hydrogen peroxide (H2O2), partly due to SPCH-driven repression of H_2_O_2_-scavenging enzymes CAT2 and APX1. As hydrogen peroxide is also a byproduct of oxidative phosphorylation (Wong et al., 2017), future work should explore a potential link between oxidative phosphorylation and stomatal development.

Manipulations of starting meristemoid size or growth rates (in tetraploid and *ctr1* plants, respectively) affect the rate of amplifying divisions and thus the stomatal index of mature leaves (Figure 4). In general, coupling cell size to differentiation may provide a quantitative tuning point to integrate internal and external signals that control meristemoid self-renewal. Cell size may be an especially attractive integrator for several reasons. First, in principle, meristemoid shrinkage can be tuned to a wide range of values, allowing flexible control of the number of divisions before differentiation. Second, many environmental and hormonal inputs are known to influence cell size in plants by controlling the rate of cell expansion and/or cell cycle length (Gonzalez et al., 2012; Moreno et al., 2020; Vaseva et al., 2018). Lastly, the evolution of tissue- and stage-specific cell cycle regulators (Han et al., 2021; Han & Torii, 2019) may allow organisms to fine-tune cell size in specific cell lineages or developmental stages.

Over the years, cell biologists have made significant inroads on the question of why cell size matters. A growing body of work has shown that size matters for division competence (Xie et al., 2022), biosynthetic capacity (Schmoller & Skotheim, 2015), metabolic flux (Homem et al., 2014), and stem cell exhaustion (Lengefeld et al., 2021). Our study adds fate specification to the compendium of size-regulated processes, building on previous work in *C. elegans* (Hubatsch et al., 2019) and *Volvox carteri* (Kirk et al., 1993) germ cell fate specification, and on somatic stem-cell lineages in Drosophila (Homem et al., 2014).

## Materials and methods

### Plant material and growth conditions

All *Arabidopsis* lines used in this study are in the Col-0 background, and wild type refers to this ecotype. *Arabidopsis* seeds were surface sterilized by bleach or 75% ethanol and stratified for 2 days. After stratification, seedlings were vertically grown on ½ Murashige and Skoog (MS) media with 1% agar for 3-14 days under long-day conditions (16 hours light/8 hours dark at 22°C) and moderate intensity full-spectrum light (110 μE).

Previously reported mutants and transgenic lines include: *pSPCH∷SPCH-YFP pATML1∷RCI2A-mCherry* in *spch-3* (Lopez-Anido et al., 2021), *pMUTE∷MUTE-YFP pATML1∷RCI2A-mCherry* (Davies & Bergmann, 2014), *pBRXL2∷BRXL2-YFP pATML1∷RCI2A-mCherry* (Gong, Varnau, et al., 2021; Rowe et al., 2019), *pBRXL2∷BRXL2-YFP pATML1∷RCI2A-mCherry* in *ctr1* (Gong, Alassimone, Varnau, et al., 2021), tetraploid Col-0 (Robinson et al., 2018), *p35S∷PIP2A-RFP* in *basl-2* (Rowe et al., 2019), *pBRXL2∷BRXL2-YFP pATML1∷RCI2A-mCherry* in *myoxi-i* (Muroyama et al., 2020), *pATML1∷mCherry-RCI2A pBASL∷BRX-CFP* (Rowe et al., 2019), and *pATML1∷H2B-mTFP pATML1∷mCit-RCI2A* (Robinson et al., 2018). We created *crwn1-1 pATML1∷H2B-mTFP pATML1∷mCit-RCI2A* by crossing *crwn1-1*(Dittmer et al., 2007) with *pATML1∷H2B-mTFP pATML1∷mCit-RCI2A* (Robinson et al., 2018).

### Microscopy, image acquisition, and image analysis

All fluorescence imaging, time-lapse, and time-course experiments were performed as described in (Gong, Alassimone, Muroyama, et al., 2021). To quantify SPCH protein levels (Figure 1D), we captured a time-lapse of a 3 dpg cotyledon expressing the translational reporter *pSPCH∷gSPCH-YFP*. We randomly selected meristemoids that were born during this time-lapse and recorded their subsequent behaviors (self-renewal vs. differentiation). Mean SPCH-YFP intensities were quantified as the raw integrated density of a summed projection divided by the area of the region of interest in square microns. Similarly, MUTE protein levels (Figure 1E) were measured from a time-lapse of a 3 dpg cotyledon expressing *pMUTE∷MUTE-YFP*. We segmented, tracked, and measured the mean fluorescence intensity of MUTE-YFP using the TrackMate Fiji plugin (Tinevez et al., 2017). To quantify epidermal cell size at 0 dpg (Figure 4A, B), mature embryos were dissected from seeds and stained with mPS-PI staining as described previously (Truernit et al., 2008). The stained embryos were imaged using a Leica SP8 confocal microscope and MorphographX (Barbier de Reuille et al., 2015 was used to create a surface mesh containing the epidermal signal, that mesh was then segmented to quantify cell surface areas. In Figures 2, 3 and 4, meristemoid cell area was measured using the polygon tool in Fiji (description of process in Figure S1). In Figure 5, nuclear and plasma membrane signals were segmented semi-automatically using ilastik (Berg et al., 2019); nuclear and cell areas were quantified in Fiji. In Figure S6A, crescent sizes were measured using POME (Gong, Varnau, et al., 2021).

### Statistical analysis

All statistical analyses in this manuscript were performed in RStudio. Unpaired Mann-Whitney tests were conducted to compare the means of two groups using the compare_means function in the ggpubr package (Kassambara, 2020). Logistic regression was conducted with the glm.fit function with a binomial model in R (R Core Team, 2020). Classification accuracy was estimated from separate training and test datasets. Briefly, 5-fold cross validation was used to split a dataset of n = 95 cells into pairs of training and test data. In each case, the training dataset was used to estimate the logistic with the glm.fit function. Then, for each cell in the test dataset, cell size at birth was used to compute a division probability according to the logistic and a predicted behavior (division or differentiation) was assigned by binarizing the probability with threshold 0.5. Accuracy is calculated as the percentage of cells with correctly predicted behavior, and the average accuracy across all 5 cross validations was reported. For all graphs, p-values from the unpaired Mann-Whitney tests or logistic regression model were directly labeled on these graphs.

The transition size for division/differentiation was operationalized as the size at which 50% of cells differentiate. As this is conceptually equivalent to estimates of the LD50 value, the amount of a toxin that causes death in half of the subjects, we used the dose.p function from the R package MASS (Ripley, 2002) to obtain point estimates and associated error for the transition size.

### Computation models and simulations

The lineage decision tree model and all associated simulations were built and performed in MATLAB. The lineage decision tree model is a stochastic, asynchronous rule-based model of meristemoid progression through asymmetric division, differentiation, and growth. The starting sizes of 10,000 cells were randomly drawn from a gamma distribution (4, 33.3). Cells with starting sizes below 40 μm^2^ were discarded (about 5% of cells). The cells then divided asymmetrically with a division asymmetry drawn from a beta distribution (6.8, 14.6) with a noise factor ±0.05, each forming a smaller daughter cell or meristemoid and a larger daughter cell or SLGC. SLGCs were discarded, while the meristemoid differentiated with some probability based on its current size using the binomial CDF at (current size, 100, 0.32). For instance, a cell of 32 square microns would have a 50% chance of differentiating. If the cell did not differentiate, it grew by 3% ± 0.005% per hour to the power of a random cell cycle length, with the cell cycle length drawn from another gamma distribution (10, 20). Calculating growth to the power of cell cycle length allows for asynchronicity (individual cells are different ages). Parametric distributions were obtained through biological measurements. The fittings of the starting size of the meristemoid population, division asymmetry, and cell cycle length to gamma or beta distributions were conducted with the fitdist function from the fitdistrplus package (Delignette-Muller & Dutang, 2015). Cell sizes were rounded to the nearest integer μm^2^. Additional noise was introduced (+/−) to reflect uncertainty.

## Acknowledgments

We thank members of the lab and the Stanford cell size control community for engaged discussions and advice. We thank Dr. Ao Liu for experimental verification of tetraploid lines, and Dr. Andrew Muroyama for timelapses of *myoxi-i* lines.

## Contributions of authors

YG, RD, GOA and HFF contributed equally, uniquely and irreplaceably to this collaboration and should be considered first authors. YG contributed conceptualization, data acquisition, analysis and interpretation, especially in Figures 1, 2 and 4. RD created and analyzed computational models for meristemoid behavior in Figure 3. GOA and HFF contributed data acquisition, analysis and interpretation, especially in Figure 5 and S1-3, 5, 7. Writing the manuscript was a collaborative effort among YG, RD, GOA, HFF and DCB. MES imaged embryos and analyzed with MorphographX to extract cell size information.

