## Supplementary figures 1-7 for "A cell size threshold triggers commitment to stomatal fate in *Arabidopsis*"

**Supplementary Figure 1. Measuring cell outcomes over time.**

**Supplementary Figure 2. Meristemoids decrease in size over successive divisions.**

**Supplementary Figure 3. Size-dependent differentiation is specific to the stomatal lineage.**

**Supplementary Figure 4. Size-dependent differentiation integrates multiple possible inputs.**

**Supplementary Figure 5. The meristemoid transition cell size does not set the final size of stomata.**

**Supplementary Figure 6. Effects of polarity on size-dependent differentiation.**

**Supplementary Figure 7. *CRWN1* controls nuclear size without affecting overall cell size.**

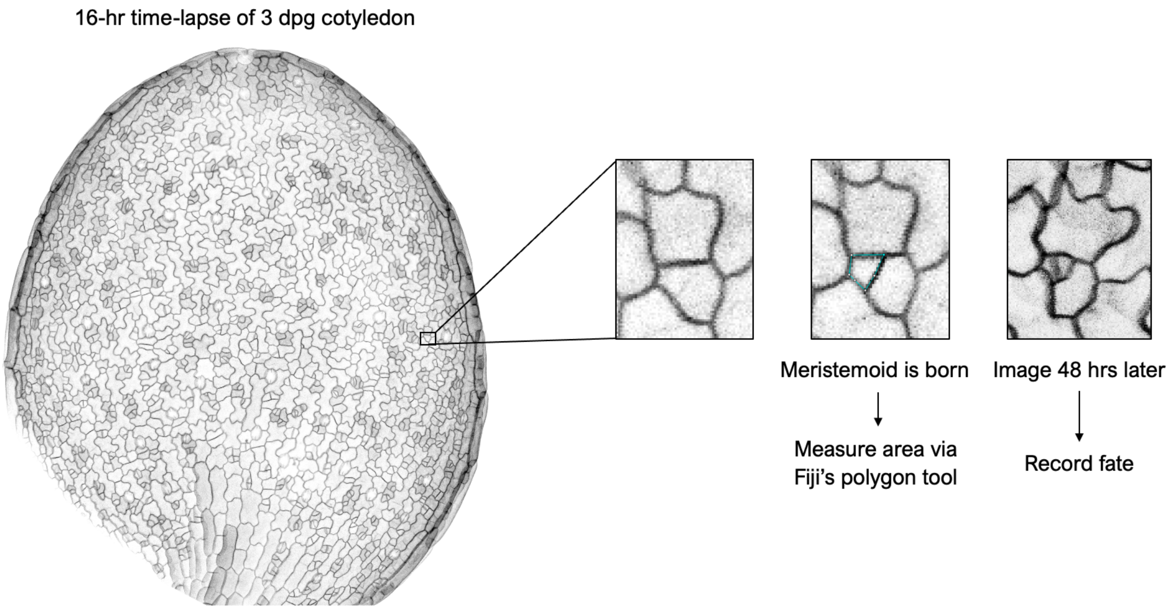


**Supplementary Figure 1. Measuring cell outcomes over time.**
Schematic of experimental setup. Whole leaf images were acquired at 40 - 60 minute intervals over 16 hours to identify cells at birth and measure key features. Then, plants were allowed to recover for 48 hours and imaged once more to record fate outcomes such as further asymmetric division (shown) or differentiation into stomata. Typical cell cycle times in this tissue are approximately 12 - 16 hours.

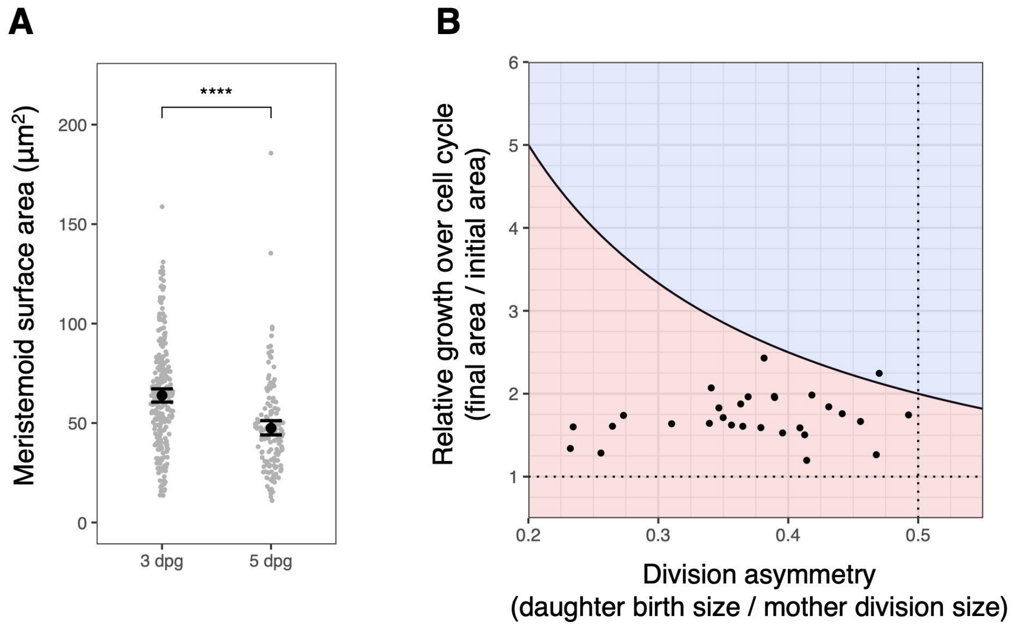


**Supplementary Figure 2. Meristemoids decrease in size over successive divisions.**
(A) Meristemoid area at 3 and 5 days post germination shows a marked population-level decrease. (B) Tracking of individual meristemoids over the whole cell cycle. Cells typically grow to less than twice their birth size (y-axis) and divide with strong physical asymmetry (x-axis), which ensures daughter cells are smaller at birth than their mothers one cell cycle earlier. The curved line marks combinations of growth and asymmetry that maintain cell size (e.g. doubling in area with 50:50 division asymmetry). P-value is calculated by Mann-Whitney test.

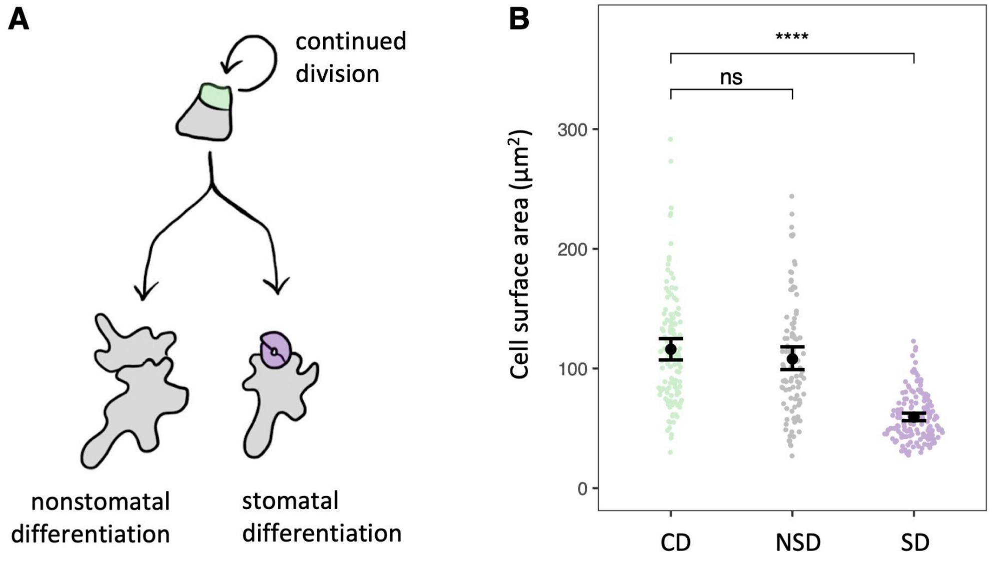


**Supplementary Figure 3. Size-dependent differentiation is specific to the stomatal lineage.**
(A) In tomato, meristemoids can differentiate into either pavement cells or stomata (Nir, Amador *et al*. 2021). (B) Nonstomatal differentiation into pavement cells (NSD) occurs at any size, but stomatal differentiation into GMCs (SD) is restricted to small meristemoids. P-values are calculated by Mann-Whitney test.

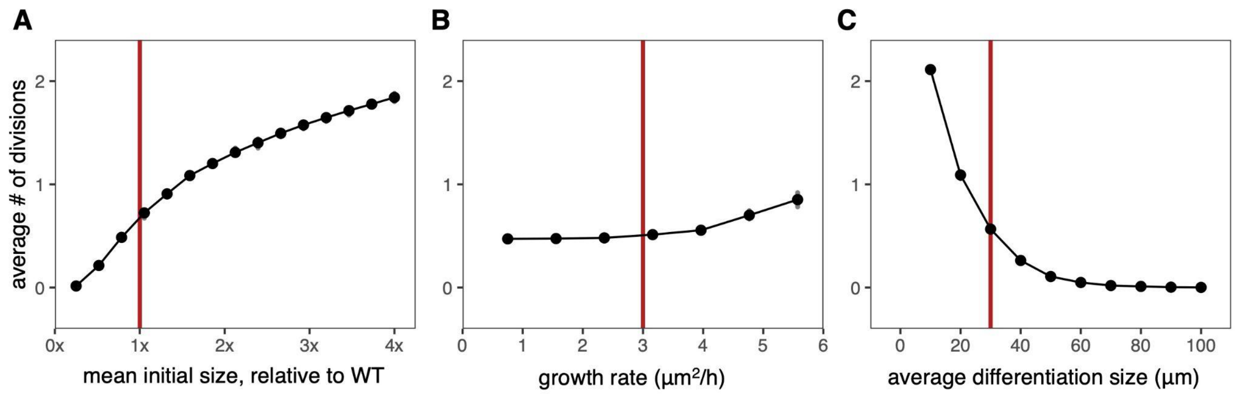


**Supplementary Figure 4. Size-dependent differentiation integrates multiple possible inputs.**
(A-C) Average number of amplifying divisions before differentiation as a function of inputs to the lineage model. (A) Initial size, in multiples of the average WT size. (B) Growth rate, in $\mu m^2$/hour. (C) Average size at differentiation. Vertical red lines indicate estimated WT values for each parameter. Black dots represent means of 1000 simulated cell lineages.

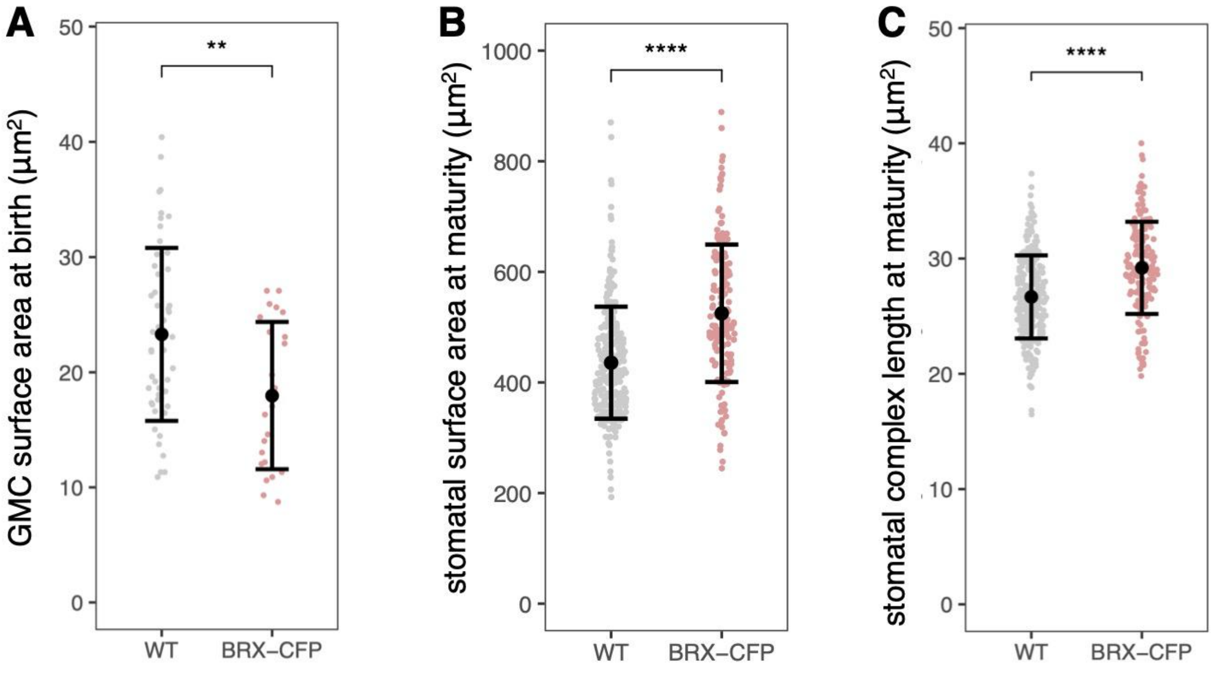


**Supplementary Figure 5. The meristemoid transition cell size does not set the final size of stomata.** (A) GMC surface area at birth in WT or a line ectopically expressing BRX in the stomatal lineage (pBASL::BRX-CFP, Rowe *et al*., 2019). (B) Surface area of mature stomatal complexes (both guard cells, measured at 21 dpg). (C) Length of mature stomatal complexes. All p-values are calculated by Mann-Whitney test.

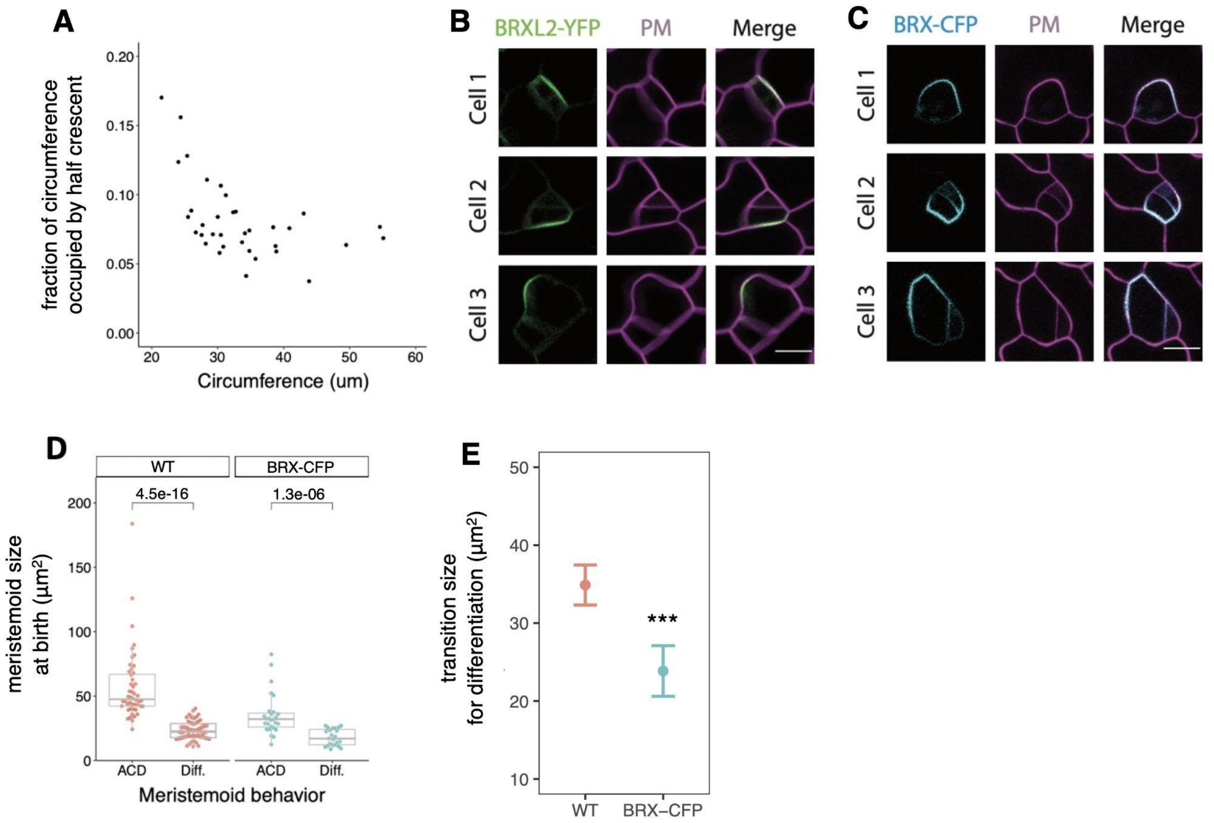


**Supplementary Figure 6. Effects of polarity on size-dependent differentiation.**

(A) Plot of the relationship between the relative proportion of plasma membrane that is occupied by BRXL2 polarity crescent (as standard deviation of the fitted curve divided by cell circumference) and the circumference of the cell (n = 35 cells). With the increase of the circumference, BRXL2 occupies a decreased proportion of the plasma membrane, indicating a lack of scaling for the BRXL2 polarity crescent. Both cell circumference and the SD of BRXL2 polar crescent are measured by POME V2.0 in FIJI. (B) Confocal images of BRXL2-YFP reporter in three representative stomatal lineage cells of different sizes. Cell 1 shows pre-divisional BRXL polarity and cells 2-3 show post-divisional BRXL2 polarity. pBRXL2::BRXL2-YFP (left), pATML1::RCI2A-mCherry (middle), and merged (right) are shown separately. (C) Confocal images of BRX-CFP reporter in three representative stomatal lineage cells of different sizes pBASL::BRX-CFP (left), propidium iodide staining (middle), and merged (right) are shown separately. Cell 1 shows pre-divisional BRXL polarity and cells 2-3 show post-divisional BRXL2 polarity. Note that in cells with particularly broad BRX crescents (e.g., Cell 2), the polar crescent can be bisected by the division plane and inherited by both daughter cells. (D) Comparison of cell size at birth between meristemoids that acquire different fates in Col-0, and the BRX-CFP reporter line (n> 50 cells/genotype). (E) Comparison of sizes at which meristemoids transition to predominantly differentiating (see methods). All p-values are calculated by Mann-Whitney test, except for (E) where a t-test was performed on the outputs from dose.p (see methods). Scale bars, 10 μm.

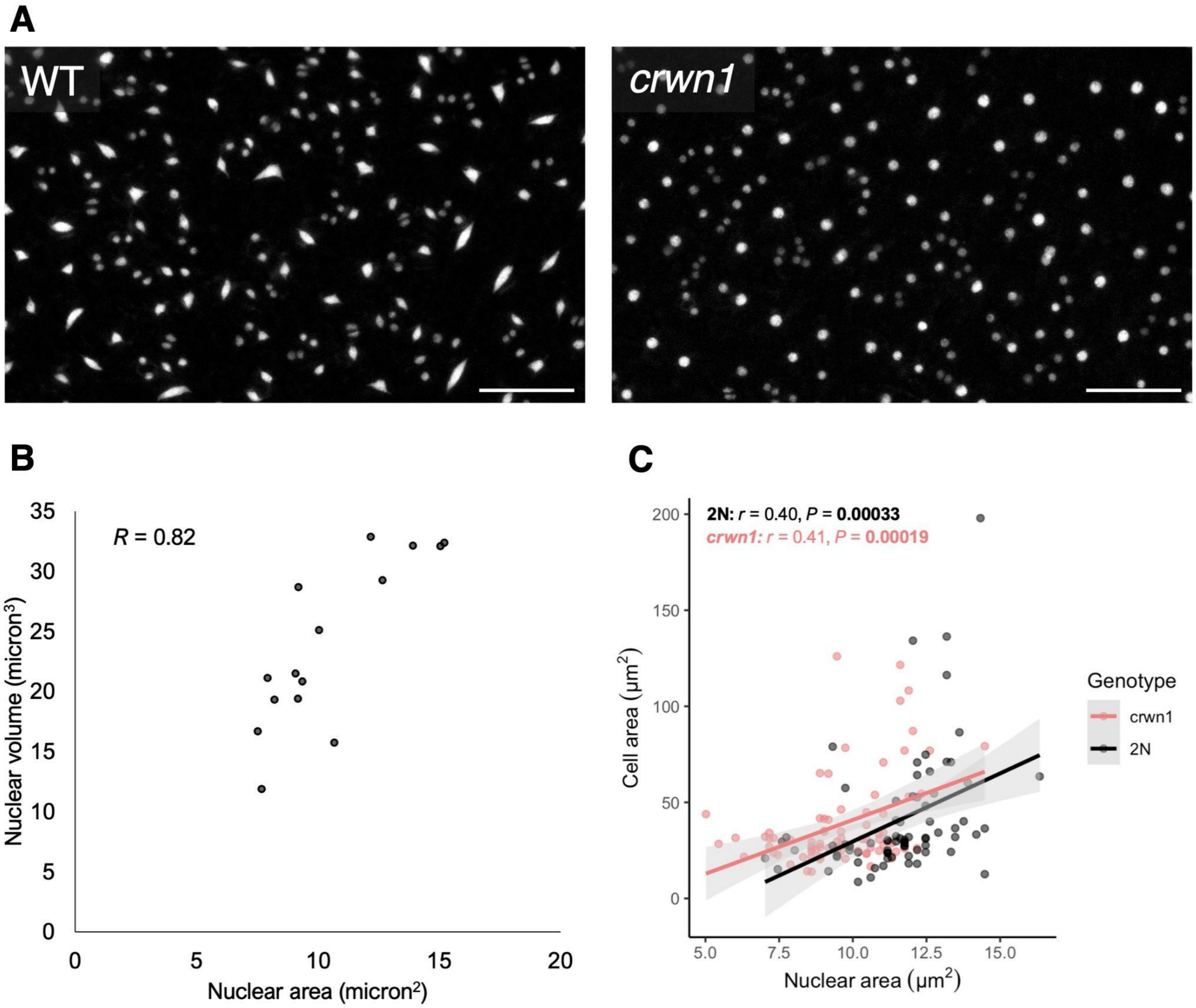


**Supplementary Figure 7.** ***CRWN1*** **controls nuclear size without affecting overall cell size.**
(A) Confocal images of WT and *crwn1* nuclei tagged with *pATML1::H2B-mTFP*. Scale bar: 50 µm. (B) Relationship between nuclear area and nuclear volume estimates. (C) Relationship between nuclear area and cell area in WT and *crwn1* plants.
